# EPH/EPHRIN SIGNALING CONTROLS PROGENITOR IDENTITIES IN THE VENTRAL SPINAL CORD

**DOI:** 10.1101/070227

**Authors:** Julien Laussu, Christophe Audouard, Anthony Kischel, Poincyane Assis-Nascimento, Nathalie Escalas, Daniel J. Liebl, Cathy Soula, Alice Davy

## Abstract

This article by Laussu et al. describes a role for Eph:ephrin signaling in controlling the identity of neural progenitors in the ventral spinal cord.

Early specification of progenitors of the ventral spinal cord involves the morphogen Sonic Hedgehog which induces distinct progenitor identities in a dose-dependent manner. Following these initial patterning events, progenitor identities have to be maintained in order to generate appropriate numbers of progeny. Here we provide evidence that communication via Eph:ephrin signaling is required to maintain progenitor identities in the ventral spinal cord. We show that ephrinB2 and ephrinB3 are expressed in restricted progenitor domains in the ventral spinal cord while several Eph receptors are more broadly expressed. Further, we provide evidence that expression of *Efnb3* and *EphA4* is controlled by Shh. Genetic loss-of-function analyses indicate that expression of ephrinB2 and ephrinB3 is required to control progenitor identities and in vitro experiments reveal that activation of Eph forward signaling in spinal progenitors up-regulates the expression of the identity transcription factor Nkx2.2. Altogether our results indicate that cell-to-cell communication is necessary to control progenitor identity in the ventral spinal cord.

## INTRODUCTION

The vertebrate neural tube is organized along its DV axis in different progenitor domains which first give rise to distinct neuronal subtypes and later on to subtypes of glial cells. Combinatorial information provided by graded Sonic Hedgehog (Shh), Wnt, BMP and FGF signaling induces the regionalized expression of homeodomain and helix-loop-helix transcription factors (TFs) that are specifically expressed in different progenitor domains (Briscoe and Novitch, 2008; Megason and McMahon, 2002). For instance, progenitors of motor neurons (pMNs) express the identity transcription factor (iTF) Olig2 while adjacent progenitors (p3) which will give rise to v3 interneurons express the iTF Nkx2.2. Ventral patterning of the spinal cord is mainly controlled by the morphogen Sonic Hedgehog (Shh) produced by the notochord and the floor plate (Balaskas et al., 2012; Ribes and Briscoe, 2009). It has been shown that in addition to doses, different exposure times to Shh also induces the expression of different sets of TFs in progenitors, thus specifying the distinct progenitor domains (Dessaud et al., 2007). Although the mode of action of Shh at the cellular level is fairly well characterized, unresolved issues in the morphogen field are to completely understand how graded information is translated into the formation of distinct domains with sharp boundaries and how progenitors maintain their identity despite varying exposure to morphogen due to tissue growth (Briscoe and Small, 2015; Rogers and Schier, 2011). One proposed mechanism is the progressive emergence of a gene regulatory network (GRN) composed of distinct iTFs whose expression is refined by cross-repressive interactions. For instance, a GRN composed of three transcription factors- Pax6, Olig2 and Nkx2.2- is required to interpret graded Shh signaling, to control the position of the boundary between the p3 and pMN progenitor domains and to provide memory of the signal (Balaskas et al., 2012; Dessaud et al., 2007). In addition, expression of these iTFs is regulated not only by the morphogen but also by other factors or signaling pathways that may affect interpretation of the morphogen gradient in a temporal manner (Rogers and Schier, 2011; Wang et al., 2011). Lastly, additional mechanisms such as cell sorting could be involved in defining and/or maintaining domain boundaries and thus indirectly participating in progenitor specification (Lei et al., 2004; Wang et al., 2011; Xiong et al., 2013).

Eph:ephrin signaling is a cell-to-cell communication pathway that has been implicated in numerous developmental processes (Kania and Klein, 2016; Lisabeth et al., 2013). One of its major biological functions is to control cell adhesion and repulsion events in developing and adult tissues thus leading to the establishment and/or maintenance of axon tracts and tissue boundaries (Cayuso et al., 2015; Fagotto et al., 2014). In addition, Eph:ephrin signaling has also been shown to control various aspects of neural progenitors development and homeostasis in the cortex including self-renewal, proliferation, quiescence and differentiation (Laussu et al., 2014). In the developing spinal cord, Eph:ephrin signaling has been prominently studied in post-mitotic neurons, specifically in axon guidance and fasciculation of motor neurons (Kao et al., 2011; Luxey et al., 2013; Luxey et al., 2015), as a consequence, virtually nothing is known on the function of this pathway in spinal progenitors. Here, we questioned the role of Eph:ephrin signaling in specifying progenitor domains in the ventral spinal cord.

## RESULTS

### Eph and ephrin expression in progenitors of the ventral spinal cord

A survey of members of the B-type Eph receptor family in the mouse ventral spinal cord (Figure 1A) indicated that spinal progenitors co-express several EphB receptors, as well as EphA4, as shown by in situ hybridization (Figure 1B-D) and immunofluorescence (Figure 1E-G). Concerning B-type ephrin ligands, in situ hybridization at different developmental stages reveals that while *Efnb1* is not expressed at significant levels in progenitors of the ventral spinal cord (Figure 1H-J), both *Efnb2* and *Efnb3* are expressed in subsets of these cells. More precisely, at all stages analyzed, *Efnb2* is expressed by progenitors located at an intermediate dorso-ventral position within the spinal cord, its expression never extending to the ventral-most region (Figure 1K-M). Conversely, expression of *Efnb3* is highest in the ventral-most region of the spinal cord at all stages analyzed, with a lower expression extending more dorsally (Figure 1N-P). Because *Efnb2* and *Efnb3* were expressed in distinct progenitor domains of the spinal cord, we asked whether these corresponded to progenitors with distinct identities, namely pMN progenitors expressing Olig2 and p3 progenitors expressing Nkx2.2 (Figure 1A). Because the expression of *Efnb2* in progenitors of the ventral neural tube was barely detectable by in situ hybridization we took advantage of a reporter mouse line that expresses H2BGFP under the control of the *Efnb2* endogenous promoter (Davy and Soriano, 2007). The benefit of this reporter strategy is that H2BGFP accumulates in the nucleus thus highlighting low domains of expression and facilitating co-expression analyses. In accordance with in situ hybridization data, H2BGFP expression was detected in a restricted population of neural progenitors from E9.5 to E11.5 (Figure 2A-D). Co-staining with Olig2 revealed that the expression domain of *Efnb2* overlapped with the Olig2^+^ (pMN) domain, however, at all stages, *Efnb2* expression extended more dorsally than Olig2 expression (Figure 2A-L). Interestingly, co-staining with Olig2 and Nkx2.2, the iTFs for pMN and p3 respectively, revealed that the ventral boundary of *Efnb2* expression corresponds to the p3/pMN boundary (Figure 2M-P). Concerning *Efnb3*, in situ hybridization followed by immunostaining for Olig2 showed that the highest domain of *Efnb3* expression corresponds to Olig2^-^ floor plate and p3 progenitors (Figure 2Q-S). Altogether, these expression analyses demonstrate that all progenitors of the ventral spinal cord co-express several Eph receptors, and that ephrinB2 and ephrinB3 are differentially expressed in pMN and p3 progenitors (Figure 2T).

**Figure 1.**
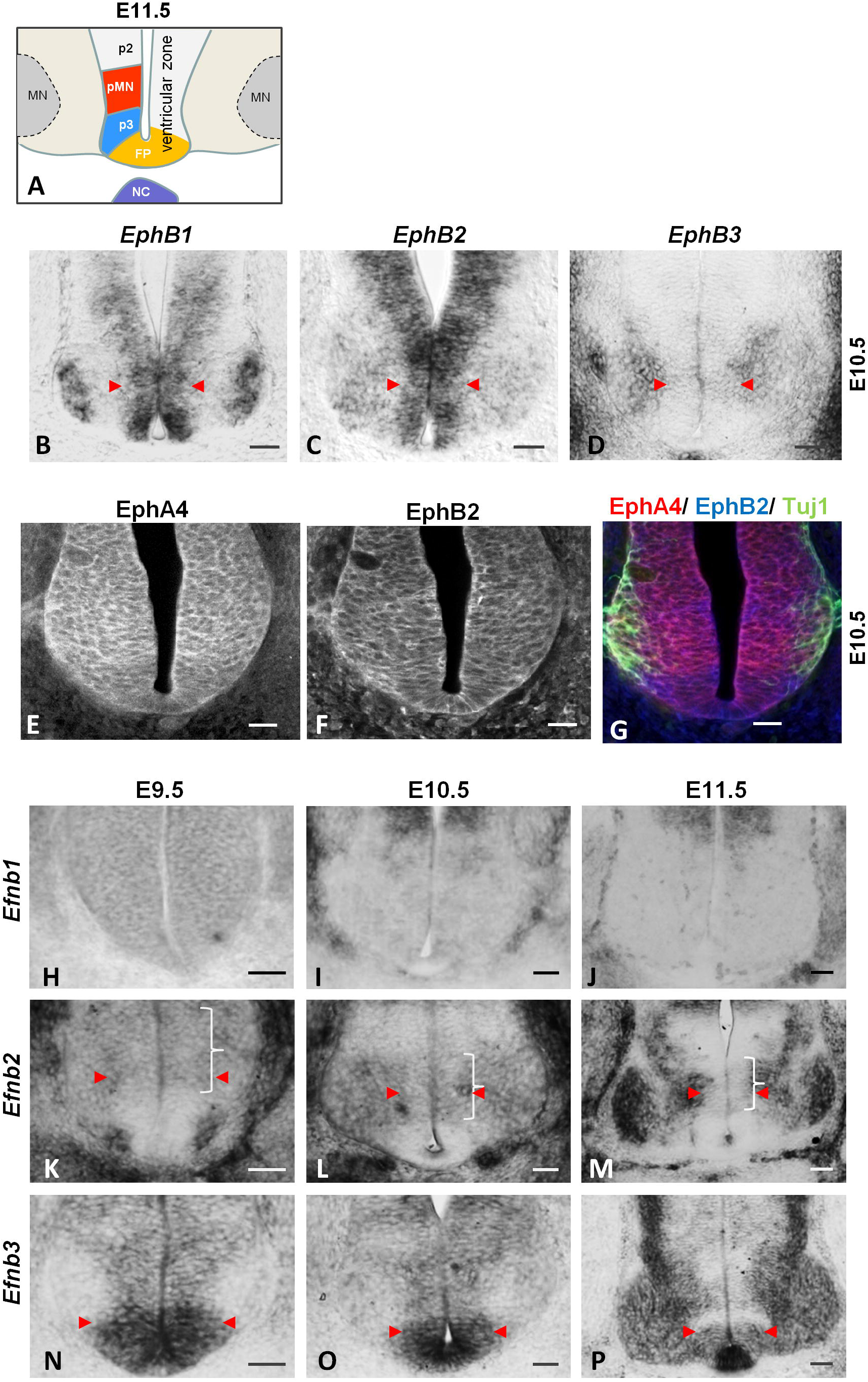
Eph receptors and ephrins are expressed in progenitors of the ventral spinal cord. A Schematic representation of the ventral spinal cord at E11.5. Progenitors are located in the ventricular zone, three progenitor domains are shown: p2, pMN and p3. Differentiated motor neurons (MN) are located laterally in the mantle zone. B-D. Expression of *EphB1* (B), *EphB2* (C) and *EphB3* (D) was monitored by in situ hybridization on transverse sections of E10.5 embryos. Scale bars: 50 μm. E-G. Transverse sections of E10.5 embryos were immunostained to detect EphA4 (E, red), EphB2 (F, blue) and differentiated neurons (Tuj1, green in G). Scale bars: 40 μm. H-P. Expression of *Efnb1* (H-J), *Efnb2* (K-M) and *Efnb3* (N-P) was monitored by in situ hybridization on transverse sections of E9.5, E10.5 and E11.5 embryos, as indicated. Scale bars: 50 μm. Arrowheads indicate the border of the ventricular zone corresponding to domains of expression. Brackets indicate domains of low expression of *Efnb2* in progenitors. FP: floor plate, NC: notochord.

**Figure 2.**
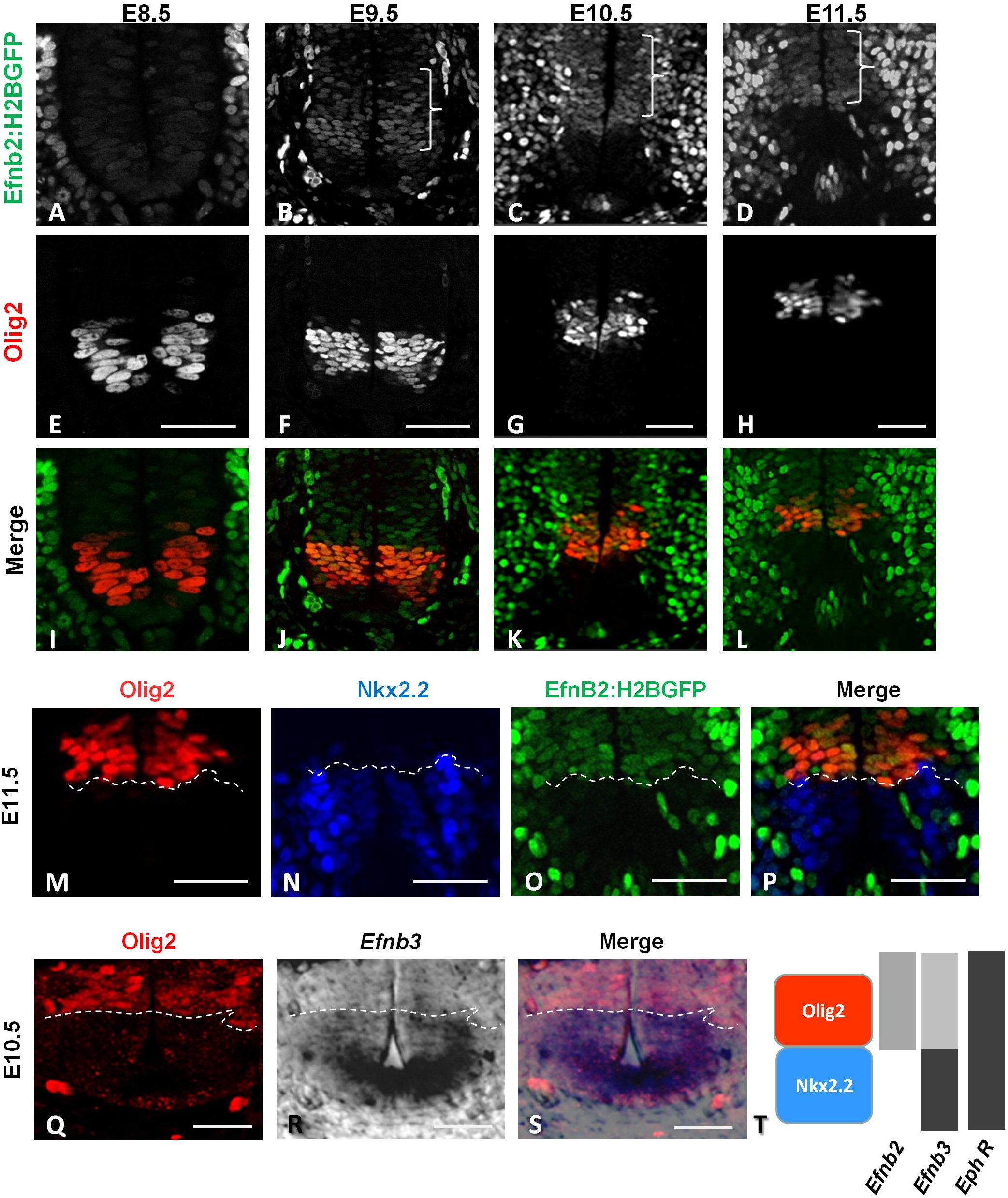
EphrinB2 and ephrinB3 are expressed in complementary domains in progenitors of the ventral spinal cord. A-L. Transverse sections of *Efnb2^+/GFP^* embryos at E8.5, E9.5, E10.5 and E11.5 (as indicated) were immunostained to detect Olig2 (E-H). Epifluorescence is shown on (A-D) and merged images are shown on (I-L). Scale bars: 50 μm. M-P. Brackets indicate domains of low expression of *Efnb2:H2BGFP* in progenitors. Transverse sections of *Efnb2^+/GFP^* E11.5 embryos were immunostained to detect Olig2 (M) and Nkx2.2 (N). Epifluorescence is shown on (O) and a merged image is shown on (P). The dashed line marks the p3/pMN boundary. Scale bars: 50 μm. Q-S. Transverse sections of wild type E10.5 embryos were processed first for *Efnb3* in situ hybridization (R) and second for Olig2 immunostaining (Q). A merged image is shown on (S). The dashed line marks the p3/pMN boundary. Scale bars: 25 μm. T. Schematic representation of *Efnb2*, *Efnb3* and Eph receptors (Eph R) expression in relation to pMN and p3 progenitors domains.

### Expression of *Efnb3 and EphA4* is regulated by Shh signaling

Shh is the main actor of ventral patterning in the spinal cord, raising the possibility that it could be involved in setting up Eph and ephrin expression domains in the ventral spinal cord. To address this we developed an in vitro culture system in which primary spinal progenitors (SPs) are grown as neurospheres. In basal, growing conditions, very few cells express neuronal (Tuj1) or astrocyte (GFAP) markers (Figure 3A), indicating that the majority of these cells are progenitors. Scattered expression of markers of differentiated motor neurons (Foxp1, Islet1/2) (Figure 3B) suggest that some of these progenitors are pMN. Indeed, Olig2 expression was detected in a large fraction of these cells while expression of Nkx2.2 is more restricted (Figure 3C). To characterize these cells further, we performed qRT-PCR analyses to monitor expression of members of the Eph:ephrin family, of genes involved in the Shh transduction cascade and of iTFs required for p3 and pMN specification. Both *Efnb2* and *Efnb3* are expressed at low levels in SPs while *Efnb1* and *EphA4* are highly expressed and *EphB2* is undetected (Figure 3D). Interestingly, the three iTFs forming the GRN required for p3 and pMN specification, namely *Pax6*, *Olig2* and *Nkx2*.2, are expressed in SPs (Figure 3D), in proportions consistent with immunofluorescence data. Lastly, components of the Shh signaling pathway *Ptch1* and *Gli3* are expressed at high levels in these cells while expression of *Gli1* is undetected (Figure 3D). These results show that expression of several genes expressed in progenitors of the ventral spinal cord is maintained in SPs. Next, we treated these cells with purmorphamine (an agonist of Smo which activates the Shh transduction cascade) for 24h and performed qRT-PCR analyses for several genes of interest. As expected, expression of *Ptch1* (which we used as a positive control since it is a Shh-responsive gene) was increased in response to high doses of purmorphamine, (Figure 3E). Interestingly, expression of *Efnb3* and *EphA4* was also increased in response to high doses of purmorphamine, while no change in the expression of *Efnb1*, *Efnb2* or *Pax6* was observed (Figure 3E). These results show that Shh signaling is able to regulate the expression of *Efnb3* and *EphA4* in spinal progenitors and suggest that Shh could participate in establishing their expression patterns in vivo.

**Figure 3.**
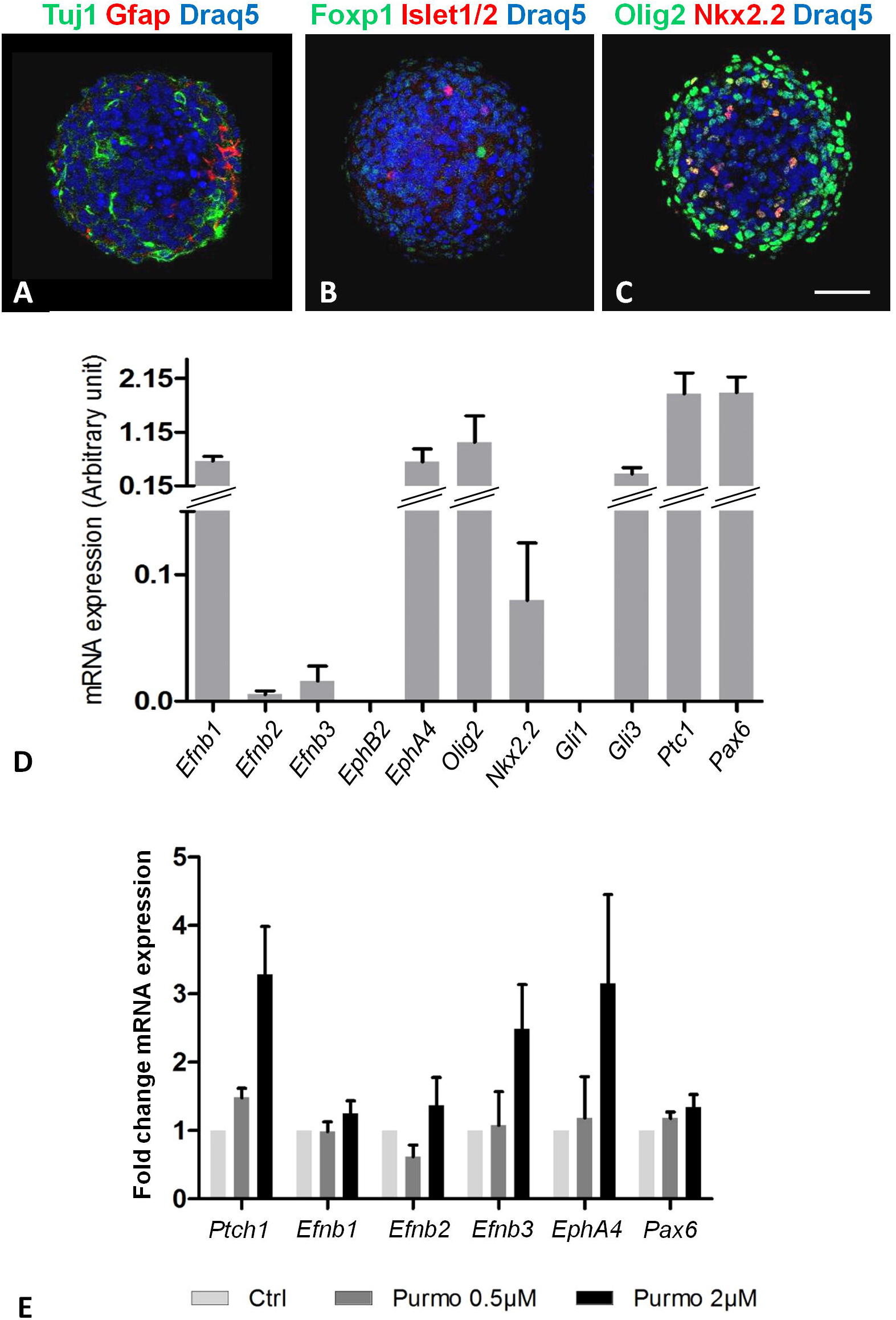
Expression of *EfnB3* and *EphA4* is up-regulated by Shh signaling. A-C. Spinal progenitors grown as neurospheres were immunostained with Tuj1 (neurons, green in A) and GFAP (astrocytes, red in A); Islet 1/2 (MN, red in B); Foxp1 (LMC MN, green in B); Olig2 (pMN and oligodendrocyte progenitors, green in C) and Nkx2.2 (p3 progenitors, red in C). Nuclei are stained with Draq5 (blue, A-C). Scale bars: 50 μm. D. Expression levels of various genes (indicated) in spinal progenitors were analyzed by qRT-PCR. The graph is representative of three independent primary cultures. E. Spinal progenitors were incubated for 24 h either with DMSO or with two doses of purmorphamine (as indicated). Expression levels of different genes (indicated) was analyzed by qRT-PCR. The graph shows fold change in expression levels for each gene in purmorphamine-treated cells compared to the control condition (ctrl). Error bars indicate s.e.m.; n=3 experiments from 3 independent primary cultures. Because of the variability in the fold change between experiments, none of the differences were statistically significant using an unpaired two-sample *t*-tests. The data should thus be interpreted as a trend.

### Loss of ephrinB3 leads to intermingling of p3 and pMN progenitors

One of the main role of Shh in the ventral spinal cord is to specify the identity of ventral progenitors, specifically pMN and p3 progenitors, but it has also been proposed to play a role in sorting pMN and p3 progenitors in their respective domains (Lei et al., 2004; Wang et al., 2011; Wijgerde et al., 2013). To ask whether Eph: ephrin signaling could play a role in these processes, we generated *Efnb2* and *Efnb3* mutant embryos. *Efnb2^-/-^* embryos exhibit precocious lethality (E10.5) due to cardiovascular defects (Adams et al., 1998; Wang et al., 1998) prompting us to generate *Efnb2* conditional mutant embryos to analyze later stages of development. We used the *Olig2-Cre* mouse line to excise *Efnb2* specifically in the motor neuron lineage, starting in pMN progenitors from E9.5 (*Efnb2^lox/lox^; Olig2-Cre* thereafter called cKO). *Efnb3* knock-out embryos (*Efnb3^-/-^* thereafter called KO) are viable and were used for all our analyses. One of the well characterized function of Eph: ephrin signaling is to maintain developmental boundaries, we thus first assessed maintenance of the p3/pMN boundary. We performed immunostaining and examined the distribution of Olig2^+^ and Nkx2.2^+^ cells in E11.5 *Efnb2* cKO and *Efnb3* KO. While the distribution of Olig2^+^ and Nkx2.2^+^ progenitors appeared similar in *Efnb2* cKO and WT embryos (Figure 4A, B), we could observe mis-localized Olig2+ progenitors in *Efnb3* KO (Figure 4C). To quantify this phenotype, we measured surfaces encompassing all Olig2^+^ or Nkx2.2^+^ nuclei on multiple transverse sections of WT and mutant embryos and deduced their region of overlap (Figure 4D, E). We then quantified the proportion of sections presenting an overlap and we measured the surface of overlap. In *Efnb2* cKO, no overlap between Olig2^+^ and Nkx2.2^+^ domains was detected (Figure 4F, G). In contrast, in *Efnb3* KO both the proportion of sections presenting an overlap and the surface of overlap between Olig2^+^ and Nkx2.2^+^ domains were increased (Figure 4H, I). These analyses reveal intermingling between Olig2^+^ and Nkx2.2^+^ progenitors in absence of ephrinB3.

**Figure 4.**
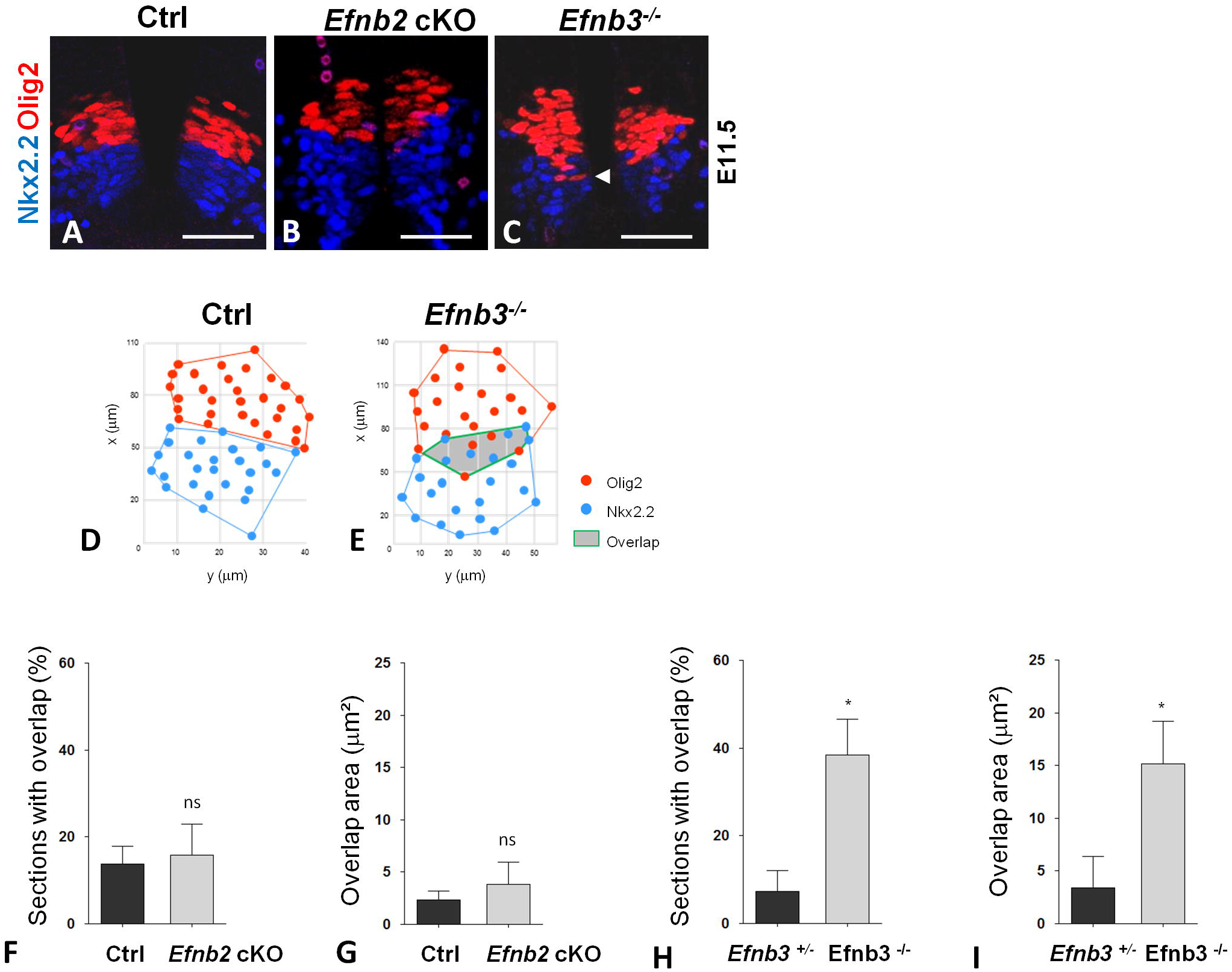
Olig2^+^ progenitors are present in the p3 domain in *Efnb3^-/-^* embryos. A-C. Transverse sections of a control (A), *Efnb2* cKO (B) and *Efnb3^-/-^* (C) E11.5 embryos were immunostained to detect Olig2 (red) and Nkx2.2 (blue). Scale bars: 50 μm. Arrowhead indicates mislocalized Olig2^+^ progenitor. D, E. Example of spatial positioning of Olig2^+^ and Nkx2.2^+^ progenitors in a non-overlap control situation (Ctrl, D) and in an overlap situation (*Efnb3^-/-^*, E). Quantification of the proportion of sections showing an overlap in control and *Efnb2* cKO embryos (F). Quantification of the surface of overlap between Olig2^+^ and Nkx2.2^+^ domains in control and *Efnb2* cKO embryos (G). Quantification of the proportion of sections showing an overlap in *Efnb3^+/-^* and *Efnb3^-/-^* embryos (H). Quantification of the surface of overlap between Olig2^+^ and Nkx2.2^+^ domains in *Efnb3^+/-^* and *Efnb3^-/-^* embryos (I). Error bars indicate s.e.m.; n=5 embryos per genotype; \**P<0.05* (unpaired two-sample *t*-tests). ns: non significant.

### EphrinB2 and ephrinB3 play opposite roles in defining the number of pMN and p3 progenitors

There are two (non exclusive) possible causes for the phenotype described above: 1) p3 and pMN progenitors are specified normally in *Efnb3* KO but the lack of ephrinB3 expression ventrally causes progenitors to intermingle; and 2) specification of p3 and pMN progenitors is altered in absence of ephrinB3 with ventral progenitors failing to acquire and/or maintain Nkx2.2 expression. To discriminate between these two possibilities, we reasoned that a specification defect may create an imbalance in p3 and pMN numbers. We thus quantified the number of Olig2^+^ and Nkx2.2^+^ spinal progenitors at two developmental stages in ephrin mutants. First we analyzed *Efnb2^-/-^* and *Efnb3^-/-^* complete knock out embryos at E9.5, a stage corresponding to initial Shh-dependent p3 and pMN patterns. No differences were observed in the distribution or number of Olig2^+^ or Nkx2.2^+^ progenitors in either mutant at E9.5 (Figure 5A-D). A small fraction of nuclei expressed both Olig2 and Nkx2.2 (progenitors of mixed identity) and this was also unchanged in absence of ephrinB2 or ephrinB3. These results show that early specification of ventral neural progenitors does not require ephrinB2 and ephrinB3 expression. Next, we quantified the number of Olig2^+^ and Nkx2.2^+^ progenitors at E11.5 in *Efnb2* cKO, *Efnb3* KO and in control embryos. Remarkably, while the total number of pMN+p3 progenitors was unchanged in *Efnb2* cKO, the ratio between pMN and p3 progenitors was modified, with a reduction in the number of Olig2^+^ progenitors balanced by an increase in the number of Nkx2.2^+^ progenitors (Figure 5E, F, I). No change in the number of progenitors with a mixed identity was observed (Figure 5I). Strickingly, *Efnb3* KO exhibit the opposite phenotype, with an increase in the number of Olig2^+^ progenitors balanced by a significant decrease in the number of Nkx2.2^+^ progenitors compared to control embryos (Figure 5G, H, J). Again, no change in the number of progenitors with a mixed identity or in the total number of p3+pMN progenitors was observed in *Efnb3* KO (Figure 5J). Importantly, using in situ hybridization, we observed no difference in *Shh* expression in *Efnb2* or *Efnb3* mutants compared to wild type embryos (Sup Figure 1).

**Figure 5.**
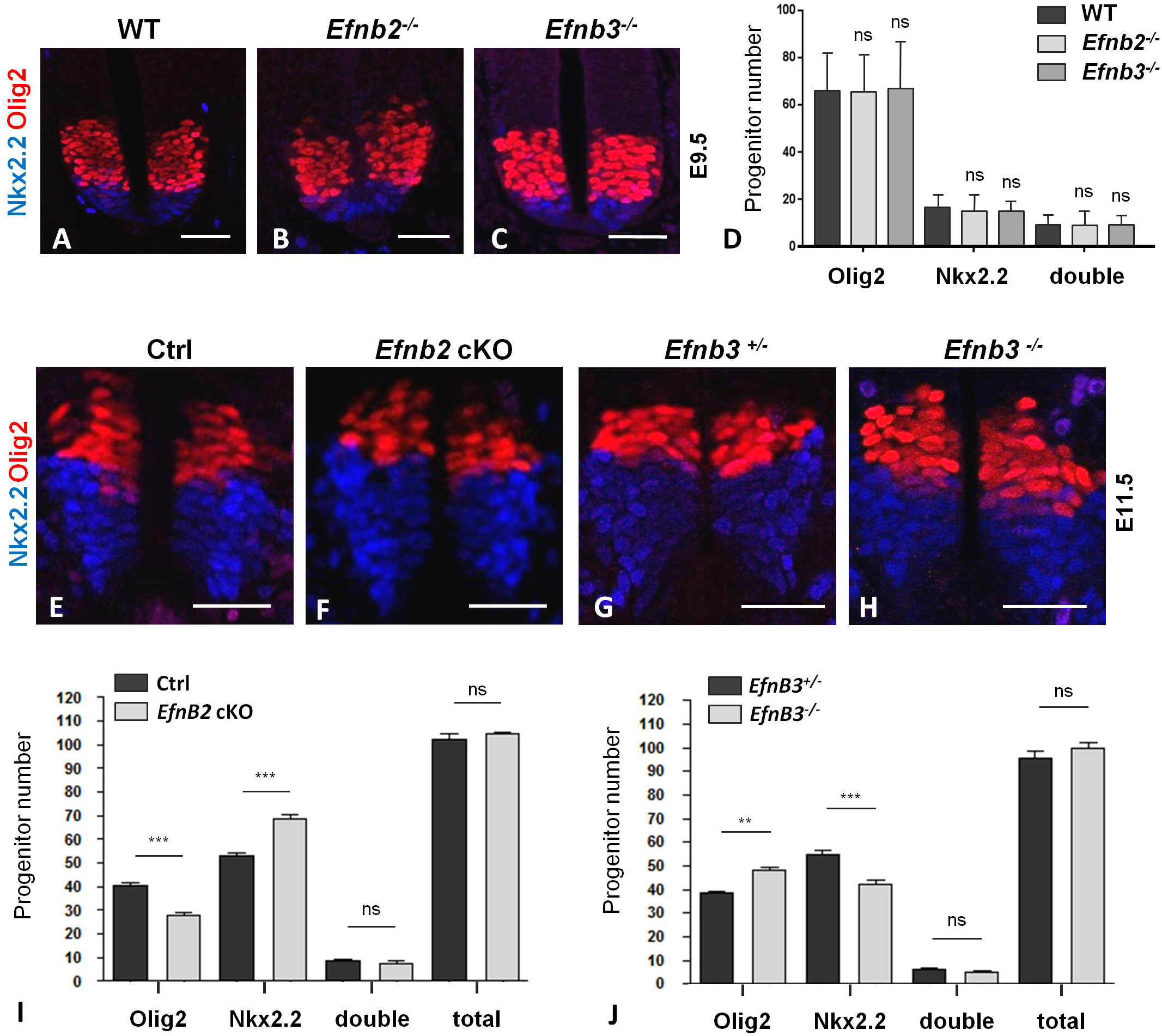
EphrinB2 and ephrinB3 play opposite roles in regulating the number of p3 and pMN progenitors. A-C. Transverse sections of wild type (A), *Efnb2^-/-^* (B) and *Efnb3^-/-^* (C) E9.5 embryos were immunostained to detect Olig2 (red) and Nkx2.2 (blue). D. The number of Olig2+, Nkx2.2+ and Olig2+/Nkx2.2+ (double) progenitors was quantified for the 3 genotypes (n=3 embryos per genotype). E-H. Transverse sections of *Efnb2^lox/GFP^* (E) *Efnb2^lox/GFP^;Olig2-Cre* (F), *Efnb3^+/-^* (G) and *Efnb3^-/-^* (H) E11.5 embryos were immunostained to detect Olig2 (red) and Nkx2.2 (blue). The number of Olig2^+^, Nkx2.2^+^ and Olig2^+^/ Nkx2.2^+^ (double) progenitors was quantified in *Efnb2^lox/GFP^* and *Efnb2^lox/GFP^; Olig2-Cre* embryos (I) and in *Efnb3^+/-^* and *Efnb3^-/-^*embryos (J) (n=5 embryos per genotype). For both graphs, total refers to the sum of Olig2^+^ and Nkx2.2^+^ progenitors. Error bars indicate s.e.m.; \*\**P*<0.01; \*\*\**P*<0.001 ns= non significant (unpaired two-sample *t*-tests).

The fact that total number of p3+pMN progenitors was unchanged suggested that rates of proliferation and differentiation of progenitors were not affected in these mutants. This was confirmed in *Efnb2* cKO by performing BrdU incorporation and immunostaining for the neuronal marker Islet1/2 which showed that proliferation and differentiation rates of Olig2^+^ progenitors were unchanged compared to wild type embryos (Sup Figure 2). Next, we assessed the number of motor neurons (pMN progeny) in *Efnb2* and *Efnb3* mutants. Despite the fact that the change in identity in *Efnb2* cKO and *Efnb3* KO affected a relatively small fraction of progenitors, this led to a change in motor neuron numbers. Indeed, *Efnb2* cKO exhibited a reduction in motor neuron numbers at later developmental stages (Figure 6A-C) while *Efnb3* KO exhibited an increase in the number of motor neurons (Figure 6E-G). In either mutant, changes in motor neuron number were not restricted to a specific motor column (Figure 6D, H). After the neuroglial transition, pMN give rise to oligodendrocyte precursors that maintain Olig2 expression and migrate in the mantle zone. In E13.5 *Efnb2* cKO, the number of Olig2^+^ cells in the mantle zone was decreased (Sup Figure 3) which is consistent with the decreased pool of Olig2^+^ progenitors in these mutants. Altogether, this data supports the interpretation that the observed changes in progenitor numbers are not due to altered differentiation rates, since progenitor and progeny numbers remain proportional in *Efnb2* and *Efnb3* mutant embryos.

**Figure 6.**
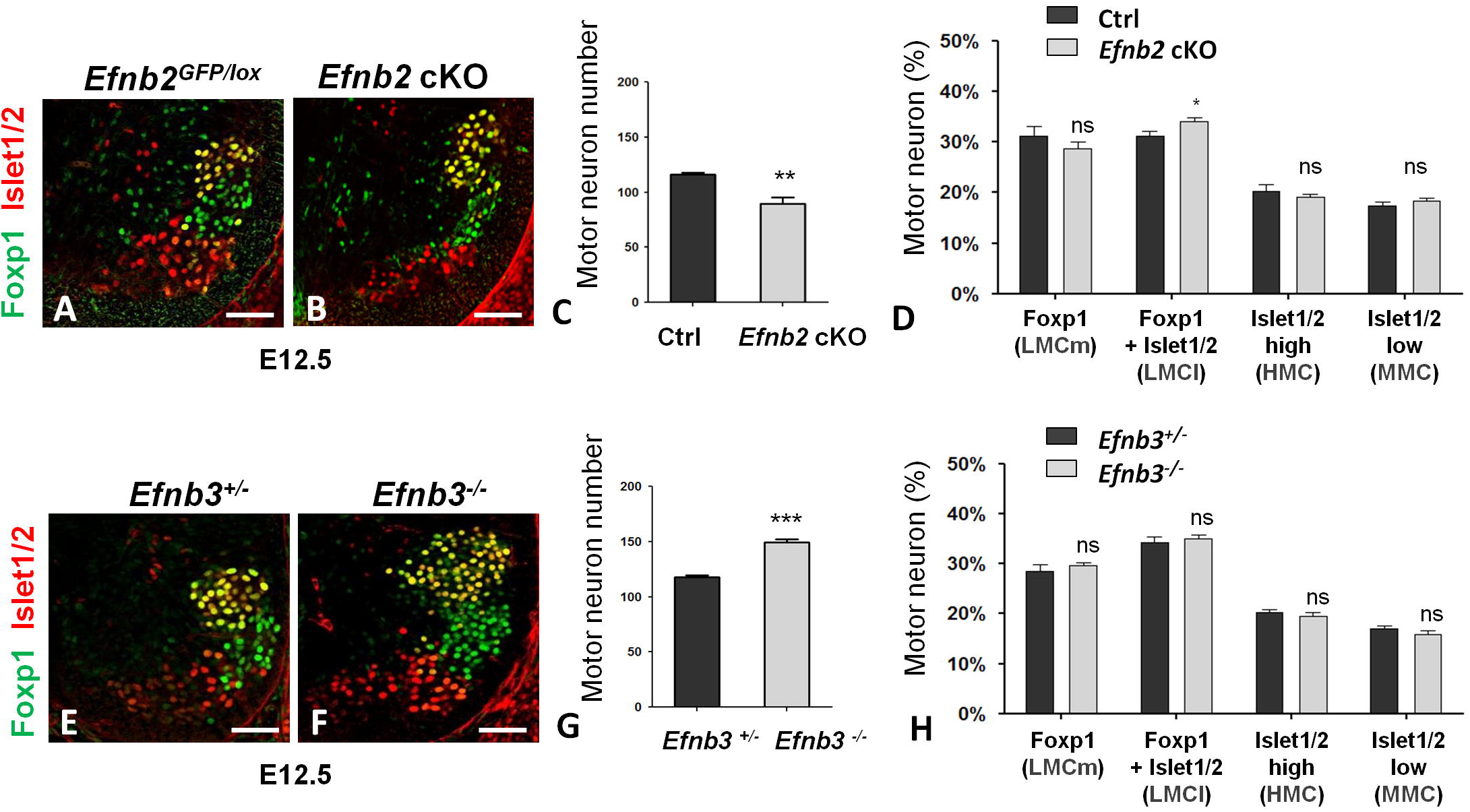
EphrinB2 and ephrinB3 inversely control motor neuron numbers. A, B. Transverse sections of E12.5 *Efnb2^lox/GFP^* (A) and *Efnb2^lox/GFP^; Olig2-Cre* (B) embryos were immunostained to detect Foxp1 (green) and Islet 1/2 (red). C. Quantification of the total number of motor neurons (Foxp1^+^ and Islet 1/2^+^) in both genotypes. D. Repartition of motor neurons in motor columns in both genotypes. E, F. Transverse sections of E12.5 *Efnb3^+/-^* (E) and *Efnb3^-/-^* (F) embryos were immunostained to detect Foxp1 (green) and Islet 1/2 (red). G. Quantification of the total number of motor neurons (Foxp1^+^ and Islet 1/2^+^) cells in both genotypes. H. Repartition of motor neurons in motor columns in both genotypes. Error bars indicate s.e.m.; n=6 embryos per group; \**P*<0.05; \*\**P*<0.01; \*\*\**P*<0.001; ns= non significant (unpaired two-sample *t*-tests). Scale bar: 50 mm.

Collectively, these results indicate that ephrinB2 and ephrinB3, which are differentially expressed in pMN and p3 progenitors, are required to maintain these respective progenitor identities after initial specification events.

### Eph forward signaling controls iTF expression

To decipher the role of Eph:ephrin signaling in specification events, we used primary cells isolated from spinal cord explants and cultured for 24 h in various conditions. As a control for this experimental set up, we treated primary cells with purmorphamine to activate Shh signaling and performed immunostaining for Nkx2.2 and Olig2 (Figure 7A-C). Consistent with the known role of Shh in promoting the p3 identity (Dessaud et al., 2007), purmorphamine treatment led to a dose-response increase in the fraction of Nkx2.2^+^ cells (Figure 7D). This experimental set up was thus used to assess whether activation of Eph:ephrin signaling promotes a specific identity. Remarkably, incubation of primary cells with clustered ephrinB3-Fc for 24 h to activate Eph forward signaling led to an increase in the fraction of Nkx2.2^+^ cells in proportions similar to purmorphamine treatment (Figure 7D). However, co-activation of Shh and Eph signaling pathways did not lead to a further increase in the number of Nkx2.2^+^ cells (Figure 7D), suggesting that both pathways converge on a common effector. We also quantified the effect of purmorphamine or ephrinB3-Fc treatment on the number of Olig2^+^ cells, however, the data was variable in the 3 independent experiments (Sup Figure 4A) precluding meaningful interpretation. Of note, very few cells were double positive (Olig2^+^/ Nkx2.2^+^) and their numbers did not change with any of the treatments (data not shown). Shh controls progenitor identities by regulating the expression of iTFs, including *Nkx2.2* which is one of its direct target genes. To assess short-term changes in gene expression, we performed qRT-PCR analyses on SPs treated for only 2 h with two doses of purmorphamine. This led to an increase in *Nkx2.2*, *Olig2*, *Gli3* and *Ptch1* expression, ranging from ~1.5 to 3 folds depending on the gene (Sup Figure 5). Interestingly, activation of Eph signaling with ephrinB3-Fc for this short 2 h time window specifically up-regulated the expression of *Nkx2.2* (in similar proportion to purmorphamine) while the expression of *Olig2*, *Gli3* and *Ptch1* was unchanged (Figure 7E). In contrast, incubation of SPs in clustered EphB2-Fc did not lead to a change in *Nkx2.2* expression (Figure 7F), validating the specificity of the response to clustered ephrinB3-Fc. Together, these results indicate that similar to Shh, activation of Eph signaling in spinal progenitors promotes the p3 identity.

**Figure 7.**
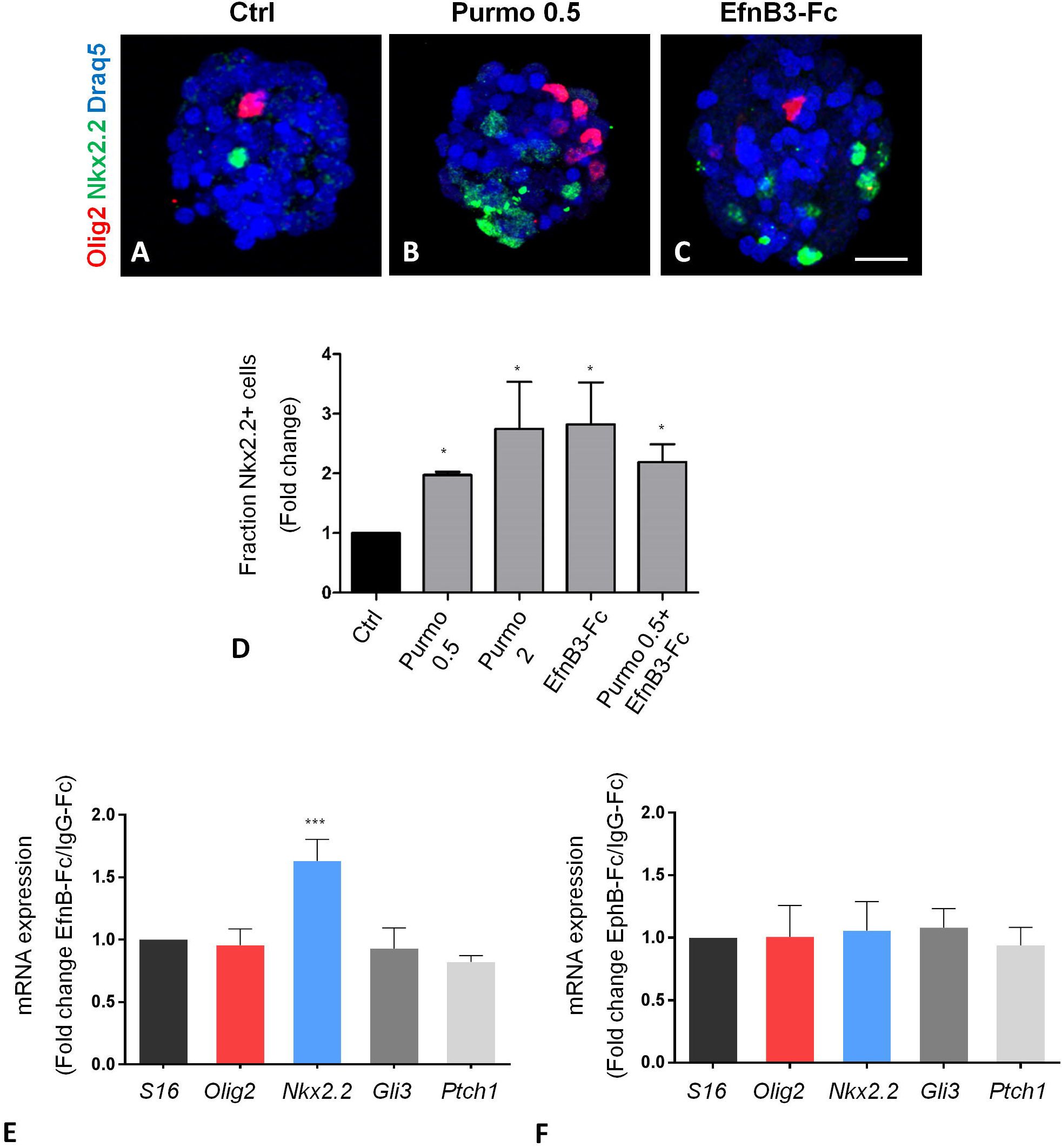
Activation of Eph forward signaling promotes the p3 fate. A-C. Representative images of primary cells isolated from E11.5 spinal cord and incubated for 24h in control medium (A), in medium containing 0.5 μM purmorphamine (B) or in medium containing 1 μg/ml ephrinB3-Fc (C). Expression of Nkx2.2 (green) and Olig2 (ref) was detected by immunostaining. Nuclei were stained with Draq5 (blue). Scale bar: 25 μm. D. Quantification of the proportion of Nkx2.2^+^ cells in control and treated conditions (as indicated). The graph represents fold change (treated vs. ctrl). Error bars indicate s.e.m.; n=3 experiments from 3 independent primary cultures; \**P*<*0.05* (one-tailed Mann Whitney test). E, F. Spinal progenitors were incubated for 2 h with either pre-clustered IgG-Fc, EfnB-Fc or EphB2-Fc. Expression levels of different genes (indicated) was analyzed by qRT-PCR in all conditions. The graphs show fold change in expression levels comparing the control condition (IgG-Fc) and either EfnB-Fc (F) or EphB2-Fc (G) stimulations. Error bars indicate s.e.m.; n=5 experiments from 3 independent primary cultures; \*\**P*<*0.01;* \*\*\**P*<*0.001*; (unpaired two-sample *t*-tests).

### *Efnb2* genetically interacts with *Shh* to control pMN number

Although the above in vitro results are consistent with the phenotype observed in *Efnb3* KO (a decrease in Nkx2.2^+^ progenitors) they did not provide insights on the phenotype observed in *Efnb2* cKO (a decrease in Olig2^+^ progenitors). We reasoned that similar to ephrinB3, ephrinB2 could be involved in promoting the expression of the iTF corresponding to its domain of expression, namely Olig2. To test for this, we used a genetic approach. Indeed, previous reports have shown that continuous Shh signaling is required to maintain Olig2 expression after initial patterning (Allen et al., 2011; Allen et al., 2007; Dessaud et al., 2010), we thus used *Shh^+/-^* as a sensitized genetic background for regulators of Olig2 expression at late stages. We generated compound *Efnb2^+/-^* and *Shh^+/-^* heterozygous embryos and quantified p3 and pMN progenitors at E11.5 (Figure 8A-D). While the number of Olig2^+^ and Nkx2.2^+^ progenitors in *Efnb2^+/-^* and *Shh^+/-^* heterozygous embryos was equivalent to wild type embryos (Figure 8E), *Efnb2^+/-^*; *Shh^+/-^* trans-heterozygous embryos exhibited a phenotype similar to *Efnb2* cKO embryos, with a decreased number of Olig2^+^ progenitors compensated by Nkx2.2^+^ progenitors (Figure 8E). These results indicate that *Efnb2* and *Shh* interact genetically to maintain Olig2 expression in progenitors of the ventral neural tube.

**Figure 8.**
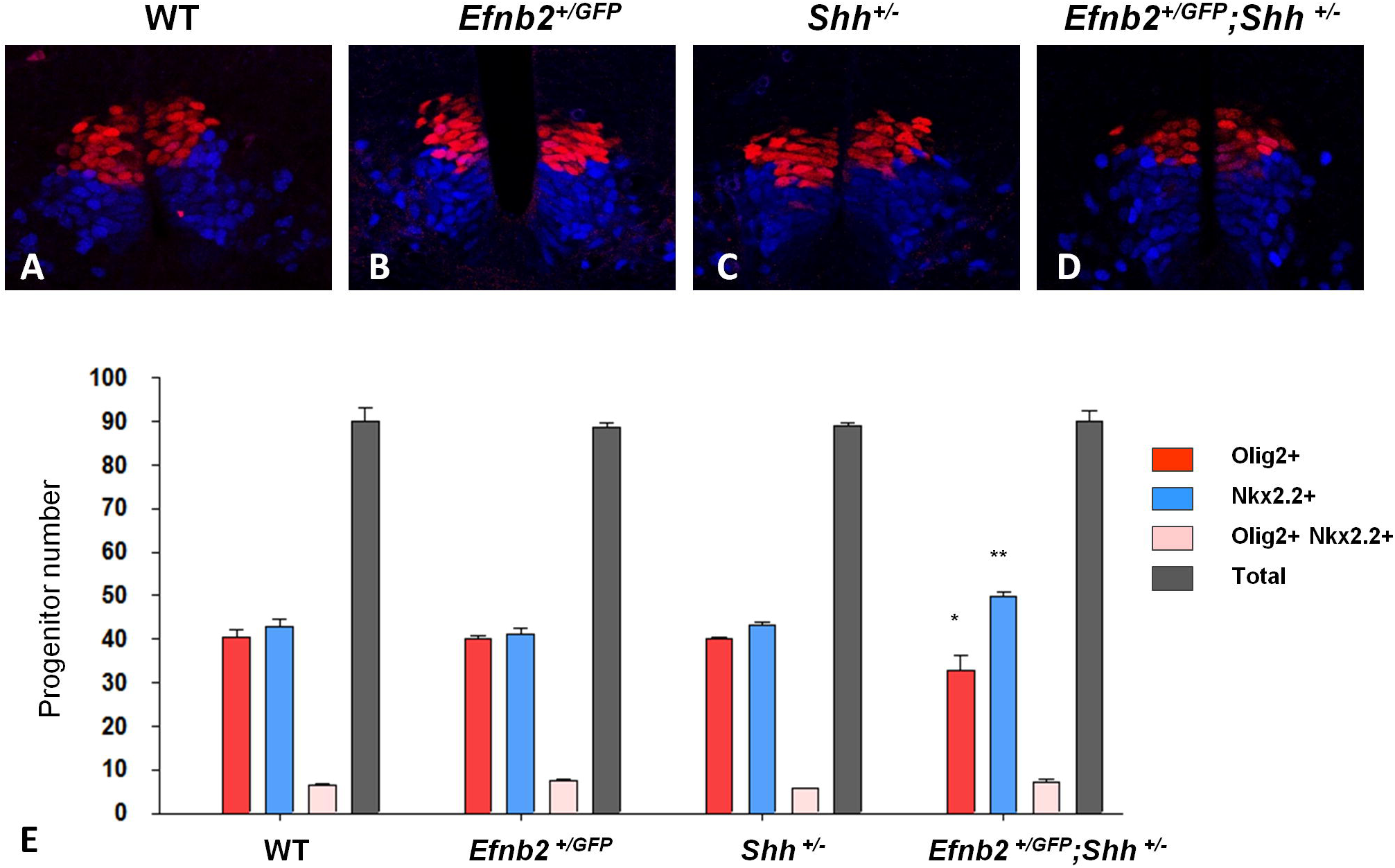
*Efnb2* and *Shh* genetically interact to control progenitor numbers. A-D. Transverse sections of E11.5 embryos of different genotypes (as indicated) were immunostained for Olig2 (red) and Nkx2.2 (blue). E. Quantification of the number of Olig2^+^, Nkx2.2^+^ and Olig2^+^/ Nkx2.2^+^ (double) progenitors was quantified for each genotype (n=5embryos per genotype). Total refers to the sum of Olig2^+^ and Nkx2.2^+^ progenitors. Error bars indicate s.e.m.; \**P<0.05*; \*\**P<0.01*; (unpaired two-sample *t*-tests).

## DISCUSSION

While early steps of ventral neural tube patterning have been extensively studied, highlighting the critical role of Shh, mechanisms that ensure fidelity in the maintenance and/or propagation of these ventral progenitor identities over time are less well characterized. Recent genetic studies indicate that continuous Shh signaling is required to maintain pMN identity since mouse mutants in which Shh signaling is altered exhibit a progressive loss of Olig2 expression (and to a lesser extend Nkx2.2 expression) (Allen et al., 2011; Dessaud et al., 2010). Here we identified Eph:ephrin signaling as a novel pathway required to maintain the identity of ventral spinal progenitors after initial patterning steps. We propose that expression of ephrinB2 and ephrinB3, respectively in pMN and p3 progenitors, is necessary to maintain the expression of Olig2 and Nkx2.2 in these cells (Figure 9). Because of the existing repressive regulatory loop between Olig2 and Nkx2.2 (Balaskas et al., 2012), modulation of Nkx2.2 expression impacts on the expression of Olig2 expression (and vice versa) thus controlling the ratio between p3 and pMN identities. Other signaling pathways have been shown to act in cooperation or in opposition to Shh to control p3 and pMN specification, including Wnt and Notch signaling (Alvarez-Medina et al., 2008; Kong et al., 2015; Robertson et al., 2004; Wang et al., 2011; Yu et al., 2008).

**Figure 9.**
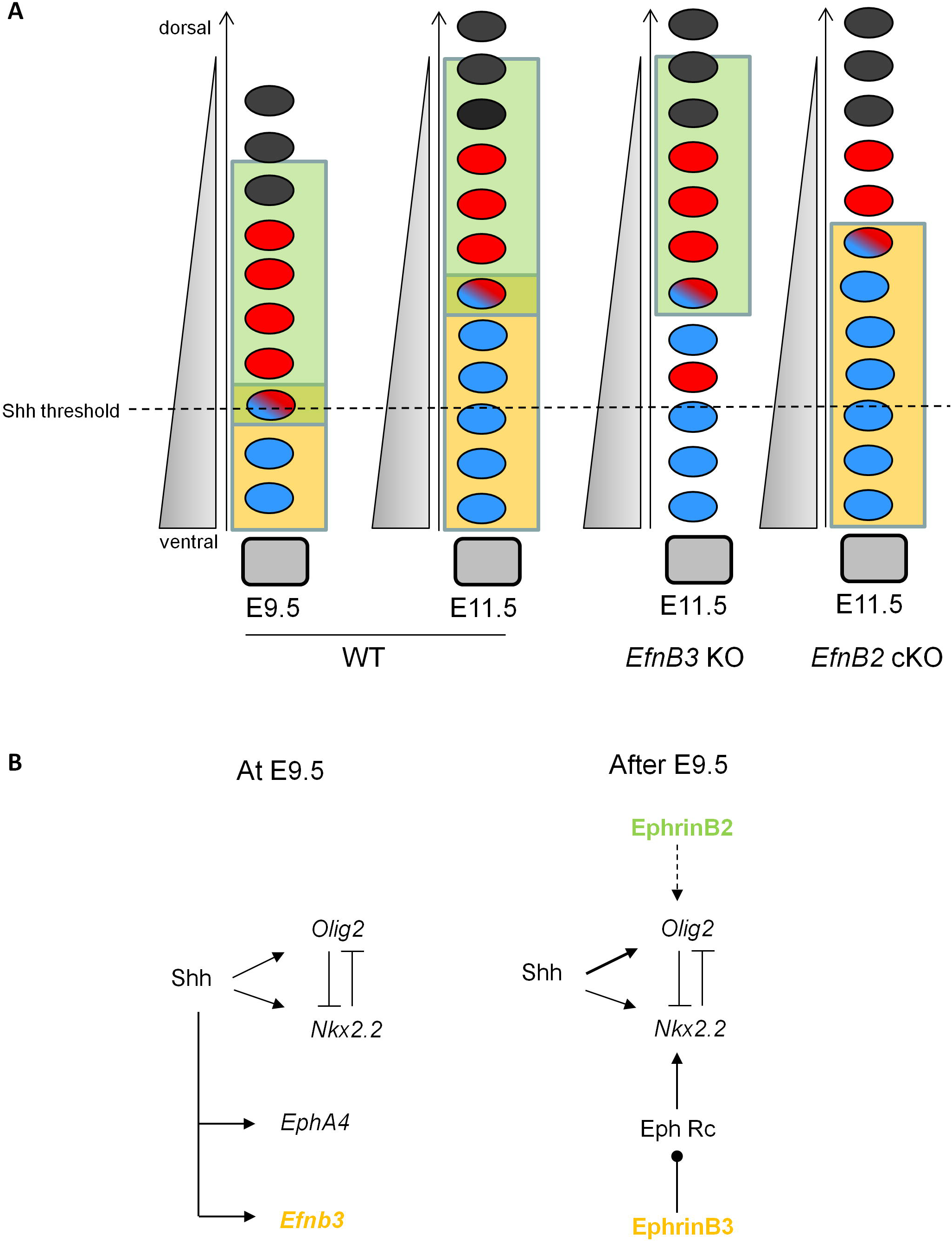
Proposed model. A Schematized representation of transverse sections of the spinal cord showing the dorso-ventral position of p3 (blue) and pMN (red) progenitors at different developmental stages and in different genetic backgrounds. Cartoons show 1) the evolution of p3 and pMN numbers over time and 2) the phenotypes observed in *Efnb2* and *Efnb3* mutants compared to WT at E11.5. Shh gradient (grey) is represented on the left hand side, while domains of ephrinB2 and ephrinB3 expression are represented in green and orange, respectively. The dotted line indicates the threshold of Shh concentration at which progenitors change identity at E9.5. At E11.5, the tissue has grown, yet p3 progenitors maintain Nkx2.2 expression despite being exposed to below-threshold doses of Shh. Nkx2.2 expression becomes progressively Shh independent while continuous Shh signaling is necessary to maintain Olig2 expression dorsally. In *Efnb3* KO, more Olig2+ progenitors are present with some detected in the p3 domain, indicating that Nkx2.2 expression is not properly maintained. In *Efnb2* cKO, pMN progenitors are fewer, suggesting a loss of Olig2 expression compensated by Nkx2.2 expression. B. At early developmental stages, Shh promotes the expression of *Olig2* and *Nkx2.2* in a dose dependent manner. Olig2 and Nkx2.2 are then locked in a mutually repressive loop. In parallel, Shh upreglates the expression of *Efnb3* and *EphA4*. At later stages, interaction between ephrinB3 and Eph receptors sets up a relay mechanism which promotes the expression of *Nkx2.2* and maintain the p3 identity. In pMN progenitors, ephrinB2 interacts with Shh to maintain the expression of *Olig2* via an unknown mechanism.

We observed that the change in identity in absence of Eph:ephrin signaling concerns only a small fraction of p3 and pMN progenitors, the majority of which maintain a correct identity in ephrin mutants. This is consistent with the mild increase in the size of the Nkx2.2^+^ domain observed in *Olig2^-/-^* embryos (Balaskas et al., 2012), suggesting that only a small fraction of progenitors adopt a p3 fate even in complete absence of Olig2. Similarly, *Tcf3^-/-^;Tcf4^-/-^* double mutants exhibit a small shift in the ratio between Nkx2.2^+^ and Olig2^+^ progenitors and this was linked to the role of Tcf3/4 in inhibiting Nkx2.2 expression in progenitors fated to become pMN (Wang et al., 2011). Using lineage tracing, it was proposed in the same study that the majority of pMN progenitors derive from cells that transiently activate Nkx2.2 expression. One attractive possibility is thus that cells of mixed identity, which represent a small fraction of p3+pMN progenitors at all stages analyzed, could be the target population, susceptible to adopt one or the other identity depending on external signals. Because of the repressive regulatory loop between Olig2 and Nkx2.2, a minor shift in expression of one of these iTFs would be amplified and result in commitment to a specific fate.

Traditionally, the role of Eph:ephrin signaling in specification processes has been linked to its function in boundary maintenance. For instance, a recent study has shown that loss of ephrinB2 in the developing cochlea leads to a switch in cell identity from supporting cell to hair cell fate and this was attributed to the mis-positioning of supporting cells into the hair cell layer (Defourny et al., 2015). Our results are not consistent with this interpretation since changes in the ratio between p3 and pMN progenitors observed in *Efnb2* and *Efnb3* mutants is opposite what would be expected from mis-positioning of progenitors. Instead, our data suggest that Eph signaling regulates specification by directly controlling the expression of iTFs. Indeed, our in vitro data shows that activation of Eph forward signaling by ephrinB3 leads to an increase in the expression of *Nkx2.2* and in the number of Nkx2.2^+^ cells which is consistent with the reduction in the number of Nkx2.2^+^ cells observed in *Efnb3* KO. Although we cannot rule out that ephrinB3 also plays a role in maintaining the p3/pMN boundary, our data suggest that mis-positioned progenitors that were observed in *Efnb3* KO in fact represent mis-specified progenitors. This interpretation is consistent with a growing number of published studies reporting a role for Eph:ephrin signaling in lineage commitment or cell fate maintenance via the modulation of intracellular signal transduction pathways and gene expression, independently of cell sorting at boundaries (Ashton et al., 2012; Chen et al., 2016; Haupaix et al., 2013; Ottone et al., 2014; Picco et al., 2007; Stolfi et al., 2011).

Our genetic data shows that expression of ephrinB2 is required to maintain the pMN fate in two different Olig2^low^ genetic contexts (*Shh^+/-^* and *Olig2-Cre^+/-^*), yet the expression of *Olig2* was not modified in response to Eph activation in vitro, even when we used ephrinB2-Fc to stimulate Eph forward signaling (data not shown), or when we used EphB2-Fc to activate reverse signaling. To date we do not have a mechanistic explanation for how ephrinB2 promotes Olig2 expression. It may be possible that ephrinB2 acts in *cis*- instead of *trans*-interaction with Eph receptors, thus attenuating Eph forward signaling (and Nkx2.2 expression) in cells fated to become pMN. *Cis*-attenuation of Eph forward signaling by ephrinB2 was previously reported in spinal motor axon guidance (Kao and Kania, 2011). Alternatively, ephrinB2 may cell autonomously impact on Shh signal transduction cascade as was recently described for Notch (Kong et al., 2015). Genetic interaction between Shh and cell surface proteins has been reported previously and identified Gas1, Cdo and Boc as components of the Shh signaling pathway (Allen et al., 2007; Martinelli et al., 2007; Tenzen et al., 2006). Another possibility, which we do not favor because it is not consistent with our in vitro data, is that the distinct in vivo biological responses elicited by ephrinB2 and ephrinB3 are due to their ability to promote the formation of distinct Eph receptor heterodimers, as was recently shown for ephrinB1 and ephrinB2 which control migration vs. proliferation of intestinal cells, respectively (Jurek et al., 2016).

In conclusion, our study indicates that in addition to its well-known function on tissue patterning via boundary maintenance and formation, Eph forward signaling also exerts a specification function by directly regulating iTF expression. This will open the way to further discoveries on the role of Eph:ephrin signaling in refining morphogen-dependent tissue patterning.

## MATERIALS AND METHODS

### Mice

Ephrin mutant mice were maintained in a mixed background and genotyped by PCR. The mouse lines *Shh^ko^*, *Efnb3^ko^*, *Efnb2^lox^* and *Efnb2^GFP^* have been described previously (Davy and Soriano, 2007; Grunwald et al., 2004; Yokoyama et al., 2001). The *Olig2-Cre* mouse line (Dessaud et al., 2007) was maintained in a pure C57Bl6/J genetic background. For *Efnb2* cKO, control genotypes used in the study include *Efnb2^lox/lox^*, *Efnb2^lox/GFP^*, *Efnb2^+/GFP^* and *Efnb2^+/GFP^; Olig2-Cre*. For *Efnb3* KO, control genotypes are always *Efnb3^+/-^*. E0.5 is defined as the day on which a vaginal plug was detected. All animal procedures were pre-approved by the institution ethical committee (protocol number: MP/07/21/04/11).

### In Situ Hybridization

In situ hybridization was performed using standard protocols on 70μm vibratome sections at brachial level. Antisense RNA probes labeled with digoxigenin were used to detect in vivo gene expression with a 72 h incubation time.

### Immunostaining

All analyses for *Efnb2* cKO were performed on control and mutant littermates collected from at least two different litters. On the other hand, control and *Efnb3* mutant embryos were collected from independent litters. The number of embryo analyzed for each immunostaining and each developmental stage is indicated in the figure legends. To avoid bias in rostro-caudal axis, data was collected on thick vibratome sections covering the entire brachial region (600 μm). Antibody staining was performed following standard protocol on 70μm vibratome sections of mouse embryos at brachial level. For BrdU incorporation, pregnant dams were injected with BrdU (10mg/ml; 100mg/kg) with intraperitoneal injection. After 1 h, embryos were dissected in cold PBS and processed for subsequent immunostaining.

Antibodies used were: goat anti-Nkx2.2 (1/100, Santa Cruz Biotechnology); rabbit anti-Olig2 (1/1000, Sigma); mouse anti-Islet1/2, 39-4D5 (1/50, DSHB); rabbit anti-Foxp1 (1/200, Abcam), rabbit anti-P-H3 (1/1000, Millipore), rabbit anti-EphA4 (1/100, Santa Cruz Biotechnology), goat anti-EphB2 (1/50, R&D Systems), Tuj1 (1/1000, Covance). All secondary antibodies were from Jackson ImmunoResearch (1/1000).

### Image processing and quantification

Images were collected on a Leica SP5 confocal microscope or Nikon eclipse 80i microscope for *in situ* hybridization data. Cell numbers were collected blindly on 5 vibratome sections (n=25 confocal Z-sections) per embryo and at least 2000 nuclei were recorded per embryo. The number of embryo analyzed for each immunostaining and each developmental stage is indicated in the figure legends. Acquisitions of nuclei 2D positions and semi quantitative analyses of fluorescence intensity were performed using Fiji (Schindelin et al., 2012). Spatial distribution of progenitor subtypes was quantified using the R Project (http://www.r-project.org/), see Supplementary Information (Sup Code) for details on the code.

### Spinal progenitor and primary cell cultures

Spinal primary cells were obtained from spinal cord of E11.5 mouse embryos. Briefly, spinal cords were dissected in ice cold HBSS solution and transferred to pre-warmed medium (DMEM/F12 supplemented with 1.5 mM Putrescine, 5 mM Hepes, 3 mM NaHcO3, penicillin-streptomicin, 1× B27 supplement, 1× N2 supplement, 1× ITSS, 5 ng/ml FGF, 20 ng/ml EGF, 30% glucose). Single cell suspension was obtained by mechanical dissociation of pooled tissues. Following dissociation, cells were immediately incubated in various conditions: control medium (DMSO + pre-clustered IgG-Fc), purmorphamine (0.5 µM or 2 µM), 1 µg/ml pre-clustered ephrinB3-Fc. After 24h, cells were fixed, immunostained to detect Olig2 and Nkx2.2 and the fraction of Olig2^+^, Nkx2.2^+^ and double positive cells was manually counted on confocal images. Of note, cells were incubated at high density which induced the formation of aggregates within the 24h time window. These aggregates are different from neurospheres since cells are not clonally related. The experiment was repeated on three independent cell suspensions (obtained from three independent litters of embryos).

Spinal progenitor (SPs) grown as neurospheres were obtained from spinal cord of E12.5 mouse embryos. Briefly, spinal cords were dissected in ice cold HBSS solution and transferred to pre-warmed DMEM/F12. Single cell suspension was obtained by mechanical dissociation of pooled tissues. Cells were centrifuged at 1500 rpm and resuspended in pre-warmed cell culture medium freshly prepared (DMEM/F12 supplemented with 1.5 mM Putrescine, 5 mM Hepes, 3 mM NaHcO3, penicillin-streptomicin, 1× B27 supplement, 1× N2 supplement, 1× ITSS, 5 ng/ml FGF, 20 ng/ml EGF, 30% glucose). Cell culture medium was refreshed every 2-3 days by adding 10 % fresh medium. Spinal progenitor stem cells grown in suspension form “neurospheres” which were dissociated every 2 weeks using 0.25% Trypsin solution, diluted 1/3 and transferred to fresh cell culture medium.

For stimulations, neurospheres were dissociated with mild trypsine treatment and cells were incubated for 2 h at 37°C with 1 μg/ml pre-clustered IgG-Fc, ephrinB3-Fc or EphB2-Fc in full medium. RNA was extracted from cell pellets using TRI-reagent according to the manufacturer's instructions. 1 μg RNA was used for reverse transcription. Genomic DNA was degraded with 1 μl DNAse (RQ1, ROCHE) for 20 min at 37°C in 20 μl RNAse, DNAse-free water (W4502-Sigma) and the reaction stopped by adding 1 μl STOP solution under heat inactivation at 65°C for 10 min. Two μl dNTPS (10mM, Promega) and 2 μl oligdTs (100mM, idtDNA) were added for 5 min at 65°C then 8 μl 5X buffer, 2 μl Rnasin (N2511-ROCHE) and 4 μl 100mM DTT (Promega) were added for 2 min at 42°C. The mix was then divided in equal volumes in a RT negative control tube with addition of 1 μl water and in a RT positive tube with 1 μl superscript enzyme (Invitrogen) and placed at 42°C for 1 h. Reaction was stopped at 70°C for 15 min. cDNAs were diluted 20-fold and processed for quantitative PCR in triplicate. 10 μl diluted cDNAs was mixed with 10 μl premix Evagreen (BTIU31019, VWR) containing 1 μmM of each primers and PCR program run for 35 cycles on a MyiQ BioRad thermocycler. mRNA relative expression levels were calculated using the 2-ddCts method. For stimulation analyses, relative level of expression were normalized to *S16* and to the IgG-Fc condition. The experiment was done on at least three independent SP cultures.

### Statistical Analysis

For all analyses sample size was estimated empirically. Sample sizes are indicated in Figure legends and further details are provided in Sup Table 1. Statistical analyses were performed with GraphPad, using unpaired two-way Student *t*-test, Mann-Whitney test or ANOVA, depending on the data set. *P*<0.05 was considered statistically significant. *P* values provided in the Figures are for Student t-test, with the following notation: \**P*<0.05; \*\**P*<0.01; \*\*\**P*<0.001, ns= non significant. Details on statistical analyses and *P* values are provided in Sup Table 2.

## ACKNOWLEDGEMENTS

The Islet 1/2 antibody was obtained from the Developmental Studies Hybridoma Bank developed under the auspices of the NICHD and maintained by the University of Iowa, Iowa City, IA 52242. We thank Dr Novitch and Dr Jessell for sharing the *Olig2-Cre* mice and Dr Grunwald for the *Efnb2^loxlox^* mice. We are greatful to Dr Greenberg for sharing the phospho-EphA4 antibody. Dr Kania and Dr Henkemeyer provided some molecular reagents used in this study. We are grateful to Brice Ronsin for his help with confocal microscopy (TRI Imaging Core Facility) and to Marion Aguirrebengoa for her help with statistical analyses. We thank the ABC facility and ANEXPLO for housing mice. We are grateful for Sylvain Touret’s help for writing the R code. We thank Eric Agius, Serge Plaza and Alain Vincent for critical reading of the manuscript.

### COMPETING INTERESTS

The authors declare no conflict of interest.

### AUTHOR CONTRIBUTIONS

JL planned, performed and analyzed experiments, and he participated in writing the manuscript; CA, AK and NE performed and analyzed experiments; PA and DL collected and provided *Efnb3* mutant embryos and revised the manuscript; CS provided scientific input on the project and revised the manuscript; AD supervised the project, planned the experiments, analyzed the data and wrote the manuscript.

### FUNDING

Research in the Davy team is financed by the CNRS, by the Fondation ARC and by ANR (ANR-15-CE13-0010-01). JL received support from the French Ministère de l’Enseignement Supérieur et de la Recherche and from the Fondation pour la Recherche Médicale (FDT20140931010). DJL is funded by the Miami Project to Cure Paralysis and PAN by NIH/NINDS (NS089325).

## REFERENCES

Adams, R. H., Wilkinson, G. A., Weiss, C., Diella, F., Gale, N. W., Deutsch, U., Risau, W. and Klein, R. (1998). Roles of ephrinB ligands and EphB receptors in cardiovascular development: demarcation of arterial/venous domains, vascula. morphogenesi., and sprouting angiogenesis. Genes Dev. 13, 295–306.

Allen, B. L., Song, J. Y., Izzi, L., Althaus, I. W., Kang, J. S., Charron, F., Krauss, R. S. and McMahon, A. P. (2011). Overlapping roles and collective requirement for the coreceptors GAS1, CDO, and BOC in SHH pathway function. Dev Cell 20, 775–87.

Allen, B. L., Tenzen, T. and McMahon, A. P. (2007). The Hedgehog-binding proteins Gas1 and Cdo cooperate to positively regulate Shh signaling during mouse development. Genes Dev 21, 1244–57.

Alvarez-Medina, R., Cayuso, J., Okubo, T., Takada, S. and Marti, E. (2008). Wnt canonical pathway restricts graded Shh/Gli patterning activity through the regulation of Gli3 expression. Development 135, 237–47.

Ashton, R. S., Conway, A., Chinmay, P., Bergen, J., Kwang-Il, L., Shah, P., Bissell, M. and Schaffer, D. V. (2012). Astrocytes regulate adult hippocampal neurogenesis through ephrin-B signaling. Nat. Neurosci. 15, 1399–1407.

Balaskas, N., Ribeiro, A., Panovska, J., Dessaud, E., Sasai, N., Page, K. M., Briscoe, J. and Ribes, V. (2012). Gene regulatory logic for reading the Sonic Hedgehog signaling gradient in the vertebrate neural tube. Cell 148, 273–284.

Briscoe, J. and Novitch, B.G. (2008). Regulatory pathways linking progenitor patterning, cell fates and neurogenesis in the ventral neural tube. Philos Trans.R Lond B Biol Sci. 363, 57–70.

Briscoe, J. and Small, S. (2015). Morphogen rules: design principles of gradient-mediated embryo patterning. Development 142, 3996–4009.

Cayuso, J., Xu, Q. and Wilkinson, D. G. (2015). Mechanisms of boundary formation by Eph receptor and ephrin signaling. Dev Biol.

Chen, S., Bremer, A. W., Scheideler, O. J., Na, Y. S., Todhunter, M. E., Hsiao, S., Bomdica, P. R., Maharbiz, M. M., Gartner, Z. J. and Schaffer, D. V. (2016). Interrogating cellular fate decisions with high-throughput arrays of multiplexed cellular communities. Nat Commun 7, 10309.

Davy, A. and Soriano, P. (2007). Ephrin-B2 forward signaling regulates somite patterning and neural crest cell development. Dev. Biol. 304, 182–193.

Defourny, J., Mateo Sanchez, S., Schoonaert, L., Robberecht, W., Davy, A., Nguyen, L. and Malgrange, B. (2015). Cochlear supporting cell transdifferentiation and integration into hair cell layers by inhibition of ephrin-B2 signalling. Nat Commun 6, 7017.

Dessaud, E., Ribes, V., Balaskas, N., Yang, L. L., Pierani, A., Kicheva, A., Novitch, B. G., Briscoe, J. and Sasai, N. (2010). Dynamic assignment and maintenance of positional identity in the ventral neural tube by the morphogen sonic hedgehog. PLoS Biol 8, e1000382.

Dessaud, E., Yang, L. L., Hill, K., Cox, B., Ulloa, F., Ribeiro, A., Mynett, A., Novitch, B. G. and Briscoe, J. (2007). Interpretation of the sonic hedgehog morphogen gradient by a temporal adaptation mechanism. Nature 450, 717–20.

Fagotto, F., Winklbauer, R. and Rohani, N. (2014). Ephrin-Eph signaling in embryonic tissue separation. Cell Adh Migr 8, 308–26.

Grunwald, I. C., Korte, M., Adelmann, G., Plueck, A., Kullander, K., Adams, R. H., Frotscher, M., Bonhoeffer, T. and Klein, R. (2004). Hippocampal plasticity requires postsynaptic ephrinBs. Nat. Neurosci. 7, 33–40.

Haupaix, N., Stolfi, A., Sirour, C., Picco, V., Levine, M., Christiaen, L. and Yasuo, H. (2013). p120RasGAP mediates ephrin/Eph-dependent attenuation of FGF/ERK signals during cell fate specification in ascidian embryos. Development 140, 4347–4352.

Jurek, A., Genander, A., Kundu, P., Catchpole, T., He, X., Strååt, K., Sabelström, H., Xu, N.J., Pettersson, S., Henkemeyer, M. and Frisén, J. (2016) Eph receptor interclass cooperation is required for the regulation of cell proliferation. Exp. Cell Res. 348, 10–22.

Kania, A. and Klein, R. (2016). Mechanisms of ephrin-Eph signalling in development, physiology and disease. Nat Rev Mol Cell Biol.

Kao, T. J. and Kania, A. (2011). Ephrin-mediated cis-attenuation of Eph receptor signaling is essential for spinal motor axon guidance. Neuron 71, 76–91.

Kao, T. J., Law, C. and Kania, A. (2011). Eph and ephrin signaling: Lessons learned from spinal motor neurons. Semin. Cell Dev. Biol. 23, 83–91.

Kong, J. H., Yang, L., Dessaud, E., Chuang, K., Moore, D. M., Rohatgi, R., Briscoe, J. and Novitch, B. G. (2015). Notch activity modulates the responsiveness of neural progenitors to sonic hedgehog signaling. Dev Cell 33, 373–87.

Laussu, J., Khuong, A., Gautrais, J. and Davy, A. (2014). Beyond boundaries: Eph/ephrin signaling in neurogenesis. Cell Adh Migr. 8, 349–359.

Lei, Q., Zelman, A., Kuang, E., Li, S. and Matise, M. (2004). Transduction of graded Hedgehog signaling by a combination of Gli2 and Gli3 activator functions in the developing spinal cord. Development 131, 3593–3604.

Lisabeth, E. M., Falivelli, G. and Pasquale, E. B. (2013). Eph Receptor Signaling and Ephrins. Cold Spring Harb Perspect Biol. 5, a009159.

Luxey, M., Jungas, T., Laussu, J., Audouard, C., Garces, A. and Davy, A. (2013). Eph/ephrin-B1 forward signaling controls fasciculation of motor and sensory axons. Dev. Biol. 383, 264–274.

Luxey, M., Laussu, J. and Davy, A. (2015). EphrinB2 sharpens lateral motor column division in the developing spinal cord. Neural Dev 10, 25.

Martinelli, D.C. and Fan, C.M. (2007). Gas1 extends the range of Hedgehog action by facilitating its signaling. Genes Dev. 21, 1231–1243.

Megason, S. G. and McMahon, A. P. (2002). A mitogen gradient of dorsal midline Wnts organizes growth in the CNS. Development 129, 2087–98.

Ottone, C., Krusche, B., Whitby, A., Clements, M., Quadrato, G., Pitulescu, M. E., Adams, R. H. and Parrinello, S. (2014). Direct cell-cell contact with the vascular niche maintains quiescent neural stem cells. Nature cell biology 16, 1045–56.

Picco, V., Hudson, C. and Yasuo, H. (2007). Ephrin-Eph signaliing drives the asymmetric division of notochord/neural precursors in Ciona embryos. Development 134, 1491–1497.

Ribes, V. and Briscoe, J. (2009). Establishing and interpreting graded Sonic Hedgehog signaling during vertebrate neural tube patterning: the role of negative feedback. Cold Spring Harb Perspect Biol. 1, a002014.

Robertson, C. P., Braun, M. M. and Roelink, H. (2004). Sonic hedgehog patterning in chick neural plate is antagonized by a Wnt3-like signal. Dev Dyn 229, 510–9.

Rogers, K.W. and Schier, A.F. (2011). Morphogen gradients: from generation to interpretation. Annu. Rev. Cell Dev. Biol. 27, 377–407.

Schindelin, J., Arganda-Carreras, I., Frise, E., Kaynig, V., Longair, M., Pietzsch, T., Preibisch, S., Rueden, C., Saalfeld, S., Schmid, B. et al. (2012). Fiji: an open-source platform for biological-image analysis. Nat Methods 9, 676–82.

Stolfi, A., Wagner, E., Taliaferro, J. M., Chou, S. and Levine, M. (2011). Neural tube patterning by Ephrin, FGF and Notch signaling relays. Development 138, 5429–5439.

Tenzen, T., Allen, B.L., Cole, F., Kang, J.S., Krauss, R.S. McMahon, A.P. (2006). The cell surface membrane proteins Cdo and Boc are components and targets of the Hedgehog signaling pathway and feedback network in mice. Dev. Cell. 10, 647–656.

Wang, H., Lei, Q., Oosterveen, T., Ericson, J. and Matise, M. P. (2011). Tcf/Lef repressors differentially regulate Shh-Gli target gene activation thresholds to generate progenitor patterning in the developing CNS. Development 138, 3711–3721.

Wang, H. U., Chen, Z.-F. and Anderson, D. J. (1998). Molecular distinction and angiogenic interaction between embryonic arteries and veins revealed by ephrin-B2 and its receptor eph-B4. Cell 93, 741–753.

Wijgerde, M., McMahon, J.A., Rule, M. and McMahon, A.P. (2013). A direct requirement for Hedgehog signaling for normal specification of all ventral progenitor domains in the presumptive mammalian spinal cord. Genes Dev. 16, 2849–2864.

Xiong, F., Tentner, A. R., Huang, P., Gelas, A., Mosaliganti, K. R., Souhait, L., Rannou, N., Swinburne, I. A., Obholzer, N. D., Cowgill, P. D. et al. (2013). Specified neural progenitors sort to form sharp domains after noisy Shh signaling. Cell 153, 550–561.

Yokoyama, N., Romero, M. I., Cowan, C. A., Galvan, P., Helmbacher, F., Charnay, P., Parada, L. F. and Henkemeyer, M. (2001). Forward signaling mediated by ephrin-B3 prevents contralateral corticospinal axons from recrossing the spinal cord midline. Neuron 29, 85–97.

Yu, W., McDonnell, K., Taketo, M. M. and Bai, C. B. (2008). Wnt signaling determines ventral spinal cord cell fates in a time-dependent manner. Development 135, 3687–96.

